# Evaluation of intercellular and extracellular synthesis of biogenic amines by the aquatic moss *Vesicularia dubyana*

**DOI:** 10.1101/2024.07.19.604072

**Authors:** Alexander Belyshenko, Maria Morgunova, Ekaterina Malygina, Maria Dmitrieva, Natalia Imidoeva, Victoria Shelkovnikova, Tamara Telnova, Anfisa Vlasova, Tatiana Vavilina, Anastasia Tretyakova, Denis Axenov-Gribanov

**Affiliations:** Irkutsk State University, Irkutsk, Russia

**Keywords:** moss, biogenic amines, *V. dubyana*, histamine, UHPLC-MS/MS

## Abstract

Natural compounds of mosses are an understudied area of phytochemistry. Over the last 20 years, the amount of information on the chemistry of natural compounds in mosses has increased considerably, but is still underwhelming. Historically, only a few hundred natural products have been described for mosses, whereas the potential for bacteria or fungi is measured in hundreds of thousands of metabolites. Although metabolites with antimicrobial, antiviral, fungicidal, and cytotoxic activities have been discovered in mosses, only few studies have been devoted to compounds with neurogenic properties. In our study, the ability of the aquatic moss *Vesicularia dubyana* to synthesize biogenic amines intercellularly and extracellularly has been evaluated.

Aquatic moss was grown in laboratory conditions, and methanol extraction of cell biomass and moss-environment medium was conducted. The presence of biogenic amines was confirmed by UHPLC-MS-MRM analysis.

The following compounds were revealed in the cellular biomass of *V. dubyana*: histamine, kynurenine, kynurenic acid, tryptamine, and tyramine. Histamine was also detected in the moss-environment medium, indicating the extracellular synthesis of this natural compound. Methanol extracts of the studied moss showed low antibacterial and antioxidant activities. Thus, this study for the first time demonstrates biogenic amine synthesis by aquatic mosses.

## Introduction

Medicinal plants are widely used as alternative therapeutic agents for prevention or adjuvant therapy of various diseases (Sabovljević et al. 2016; Lunić et al. 2020). Although there is a great deal of data on bioactive metabolites in flowering plants, which have important medical applications, much less attention has been paid to the chemistry of the moss group members. Basically, this is due to the amount of available biomaterial, the small size of mosses, the difficulty of identification, and the lack of genomic and proteomic data (Valeeva et al. 2022; Asakawa and Ludwiczuk 2018; Medina et al. 2019).

Mosses are the oldest terrestrial cryptogamous plants and the second most diverse group of plants after flowering plants. Today, more than 23,000 species of mosses are known worldwide. Mosses include liverworts (9,000 species), hornworts (300 species), and true mosses, bryophytes (14,000 species). These higher plants are found in a wide variety of habitats and play an important role in maintaining microclimates, providing substrates, and participating in water and nutrient cycling (Bansal et al. 2024; Patiño and Vanderpoorten 2018; Minerva 2020, Asakawa and Ludwiczuk 2018; Christenhusz and Byng 2016).

However, their genetic capacity to maintain long-term viability under prolonged drought conditions and to produce a diverse range of secondary metabolites has enabled them to adapt to a variety of environmental conditions. This is what has allowed mosses to colonize a wide range of environments from Antarctic tundra to tropical forests and deserts (Horn et al. 2021, Patiño and Vanderpoorten 2018).

It is known that 136 species of mosses were frequently used in ethnomedicine in various communities across the globe: Africa, New Zealand, Turkey, Japan, European regions, Pakistan, India, and China. They were dried and homogenized, made into oils, ointments, decoctions, and used as seasonings. With a wide range of medicinal properties, they were utilized in treatment of liver and cardiovascular diseases, bruises, open wounds, snake bites, pulmonary tuberculosis, neurasthenia, fractures, burns, and other conditions (Sabovljević et al. 2016; Asakawa and Ludwiczuk 2018; Chandra et al. 2017; Bandyopadhyay and Dey 2022). Despite numerous comprehensive studies of various properties of moss secondary metabolites, only a limited number of studies have focused on compounds with neurogenic properties (Ludwiczuk and Asakawa 2020; Fukuyama and Asakawa 1991; 2001; 2020; Bandyopadhyay and Dey 2022; Lunić et al. 2022). For example, neurotrophic effects of mastigophorenes A, B, and D, produced by *Mastigophora diclados*, or acetylcholinesterase inhibition effects demonstrated for marsupellin A found in *Marsupella alpina*, etc.

The vast majority of studies have focused on terrestrial mosses as well as explants grown in *vitro* (Valeeva et al. 2022; Dziwak et al. 2022; Dague et al 2023). The model mosses employed in these studies include species *Physcomitrium patens, Sphagnum fallax, Ceratodon purpureus*, etc. To date, studies on aquatic moss species have been relatively limited and have mainly focused on their use as effective biomonitoring tools (Carrieri et al. 2022; García et al. 2023; Monaci et al. 2021; Świsłowski et al. 2023; Zotina et al. 2024). Similarly, sporadic work was conducted on the aquatic mosses *Calliergon cordifolium, Drepanocladus lycopodioides* (*Amblysteglaceae*), and *Fontinalis antipyretica*. For the aforementioned species, we analyzed the fatty acid content of total lipids, and composition of phospholipids (Asakawa et al. 2013).

One of key metabolite groups involved in regulation of living processes is represented by biogenic amines. They are known to play an important role as hormones, neuromodulators, or neurotransmitters. Biogenic amines are present in all living organisms (animals, plants, and microorganisms) as ancient components of irritability. Although the properties of these amines for plant organisms are not fully understood, it has been proven that monoamine neurotransmitters play an important role in plant life. They influence such processes as morphogenesis, flowering, ion permeability, photosynthesis, circadian rhythm, reproduction, fruit ripening, and adaptation to environmental changes (Akula and Mukherjee 2020; Roshchina 2016). There are also trace amines produced in the plant body and expressed in low concentrations. These include tryptamine, tyramine, and octopamine.

Our study was carried out on *Vesicularia dubyana*, the aquatic moss widely used in aquaristics. The objective of this study was to evaluate antioxidant and antibacterial activities of *V. dubyana* methanolic extracts and to assess the moss ability to produce biogenic amines including histamine, tryptamine, tyramine, as well as products of tryptophan metabolism such as kynurenine and kynurenic acid.

## Materials and methods

### Plant material

*Vesicularia dubyana* (V. Muller) Brotherus (1908), or Java moss, belongs to the family Hypnaceae (Hypnaceae). In its natural environment, this aquatic moss grows abundantly in regions with hot and humid climates. The plant is commonly found in India, Malaysia, Southeast Asian countries, and the island of Java. *V. dubyana* forms dark green cushions with its loose or dense foliage. This moss is a common plant in aquaculture. Mature plants form thickets that provide shelter for fish fry and serve as a substrate for fish spawning. The moss under study is easy to maintain and grows comfortably in low light with neutral pH.

### Growing of aquatic mosses

The aquatic moss of *V. dubyana* was grown under laboratory conditions at 18–23 °C. Transparent polypropylene aquaria were employed for moss cultivation. The aquaria were half filled with settled tap water that was changed every 10 days. Phytolamps (900–1300 Lux) with a photoperiod of 12/12 hours and a photosynthetic photon flux of 9 *μ*Mol/s were used for illumination. A submersible aquarium pump was used to mix the water mass in aquariua. The moss was grown for 4 weeks.

### Extraction of samples

Moss samples were extracted with 100% methanol (Vecton Ltd, Russia) in order to test their antioxidant and antibacterial activity, and to reveal the presence of biogenic amines during the target HPLC-MS analysis. Same measurements were performed for tap water in which the moss was grown (moss-environment medium). Plant samples were homogenized with the Potter glass homogenizer. The plant suspension was placed in polypropylene Eppendorf tubes in the presence of ten volumes of solvent. Thus, we used 10 ml of methanol to extract 1 g of *V. dubyana* wet weight. The extraction was conducted in a rotator at 60–80 rpm for 1 hour. The supernatant was transferred to Eppendorf tubes and stored at –20°C until analysis. Immediately prior to analysis, the samples were filtered using PTFE syringe filters (0.45μm) into HPLC vials.

In order to ascertain the presence of biogenic amines, we obtained water from the moss-environment medium and subjected it to UHPLC-MS-MRM analysis. Samples were pre-filtered with PTFE syringe filters and transferred to 2 mL Eppendorf tubes). To assess the antioxidant and antibacterial properties, 15 ml of water was collected and concentrated to 3 ml in a vacuum concentrator (VAC-24 Stegler, China) at 43°C. Samples were stored at –20°C until analysis. Immediately prior to analysis, the samples were filtered with PTFE syringe filters (0.45μm) into chromatography vials.

### Qualitative evaluation of antioxidant activity of extracts

For qualitative assessment of antioxidant activity, we used 2,2-diphenyl-1-picrylhydrazyl (DPPH) (Kat. D9132, Merck, Germany) according to the method of Kansci (2003). The experiment was conducted using 96-well plates with optically transparent bottoms. The wells of the plates were filled with 10 *μ*l of the analyzed methanol extract, or water in which the moss was grown, and 90 *μ*l of DPPH (0.04 mg/mL). Methanol was used as the solvent. Dissolved DPPH and methanol were used as negative controls. The plates were left in the dark for 30 minutes, for color development. The color change from pink to orange indicated the presence of antioxidants in a sample.

### Antimicrobial activity of moss extracts

Antimicrobial activities were evaluated using the disk diffusion method (Ruangpan and Tendencia 2004). Eighteen strains of microorganisms were selected as model test culture, namely: Gram-positive bacteria: *S. carnosus* B-8726, B. *subtilis* ATCC 66337, *B. subtilis* B-4537, *E. mundtii* B-12673, *M. luteus* B-7846, *M. smegmatis* AC-1339, *S. aureus* B-6646, *K. rhizophila* B-5389, B. *subtilis* 168. Gram-negative: *P. putida* B-4589, E. *persicina* 19328, E. *coli* B-6645, *P. aeruginosa* B-6643, E. *coli* lptD CamR, and *E. coli* tolC KanR. We also evaluated antifungal activity against *S. cerevisiae* Y-1173, *C. albicans* Y-3108, and *Candida utilis* Y-3089.

To perform the experiments, mother cultures of model microbial strains were cultivated during 24 hours according to the parameters shown in Table 1. Then 100 *μ*l of each of microbial culture was plated on solid nutrient media. Afterwards, Petri dishes were dried at room temperature in a laminar flow cabinet. For the experiment, 45 *μ*l of moss (and water) extracts was used to evaluate their antimicrobial activity. The extracts were applied to 5 mm diameter paper discs and dried at room temperature. The disks were then transferred to solid nutrient media with the prepared test cultures. The resulting Petri dishes were incubated at temperatures of 30 and 37 °C for 12–24 hours until zones of growth inhibition appeared. The size of the growth inhibition zones was measured with a ruler.

**Table 1.** Model strains used to estimate antimicrobial activity and parameters of cultivation.

| Test culture | Media | Temperature |
| --- | --- | --- |
| Gram-positive bacteria |  |  |
| *Staphylococcus carnosus* B-8726 | LB | 37 °C |
| *Bacillus subtilis* ATCC 66337 | LB | 37 °C |
| *Bacillus subtilis* B-4537 | MPA | 30 °C |
| *Enterococcus mundtii* B-12673 | LB | 37 °C |
| *Micrococcus luteus* B-7846 | YEPD | 30 °C |
| *Mycobacterium smegmatis* AC-1339 | ISP | 37 °C |
| *Staphylococcus aureus* B-6646 | MPA | 37 °C |
| *Kocuria rhizophila* B-5389 | MPA | 30 °C |
| *Bacillus subtilis* 168 | LB | 37 °C |
| Gram-negative bacteria |  |  |
| *Pseudomonas putida* B-4589 | LB | 30 °C |
| *Escherichia coli* lptD CamR | LB | 37 °C |
| *Escherichia coli* tolC KanR | LB | 37 °C |
| *Erwinia persicina* 19328 | LB | 30 °C |
| *Escherichia coli* B-6645 | MPA | 37 °C |
| *Pseudomonas aeruginosa* B-6643 | MPA | 37 °C |
| Fungi |  |  |
| *Candida albicans* Y-3108 | YEPD | 37 °C |
| *Saccharomyces cerevisiae* Y-1173 | YPD | 30 °C |
| *Candida utilis* Y-3089 | YEPD | 30 °C |

Composition of nutrient media is as follows: LB (tryptone — 10 g/L, yeast extract — 5 g/L, NaCl — 5 g/L, pH 7.5); MPA (nutritious dry broth based on enzymatic beef meat hydrolysate — 30 g/L, peptone — 9 g/L, pH 7.4); YEPD (glucose — 20 g/L, yeast extract — 5 g/L, peptone — 10 g/L, pH 6.5); ISP (peptone — 5 g/L, yeast extract — 3 g/L, malt extract — 3 g/L, glucose — 10 g/L, pH 7.2); YPD (soy peptone — 20 g/L, yeast extract — 10 g/L, sucrose — 20 g/L, pH 6.5).

### Target UHPLC-MS-MRM analysis

The presence of biogenic amines in *V. dubyana* cell biomass was determined using UHPLC-MS-MRM analysis. Chromatographic determination was performed using the UHPLC system Infinity II coupled to 6470B triple quadrupole-MS/MS (Agilent Technologies, Germany). The biogenic amines tested in this study are: histamine, kynurenine, kynurenic acid, tryptamine, and tyramine.

Chromatographic separation of samples was performed in the Infiniity Lab Poroshell 120 EC-C18 analytical column (2.1 mm × 50 mm, 1.9 *μ*m) using a mixture gradient of 0.01% formic acid in deionized water (A) and 0.01% formic acid in acetonitrile (B). The flow rate was 0.2 mL/min at 40 °C. The volume of injection was 3 *μ*L. Signal detection was performed in positive ionization mode under the following parameters: 3.5 kV capillary voltage; 300 °C carrier gas temperature; 5 l/min carrier gas flow rate; 45 psi nebulizer pressure; 250 °C desolvation temperature; 11 l/min desolvation gas (nitrogen) flow. Biogenic amines and their derivatives in the obtained extract were identified using analytical standards and a series of Multiple Reaction Monitoring (MRM) transitions during molecular fragmentation. To identify and estimate the concentration of histamine in the extract obtained from *V. dubyana* plant material, we employed the analytical standard of histamine dihydrochloride. To identify kynurenine, kynurenic acid, tryptamine, and tyramine, we used MRM-transition as a diagnostic parameter (Wallace 2018). Relevant data is presented in Table 2. Chromatographic separation of target compounds was performed according to the following gradient (time (minutes), % of B): 2 min, 10%; 3 min, 90%; 5 min, 90%; 6 min, 10 %; 8 min, 10%.

**Table 2.** MRM parameters of tested biogenic amines.

| No. | Metabolite | Precursor ion, Da<br>(ionization) | Product ion, Da | Collision energy<br>(V) |
| --- | --- | --- | --- | --- |
| 1 | Histamine | 112.1 (+) | 95.1 | 17 |
|  |  |  | 83.0 | 20 |
|  |  |  | 68.0 | 20 |
| 2 | Kinurenine | 209.0 (+) | 192.0 | 10 |
|  |  |  | 146.0 | 15 |
|  |  |  | 94.0 | 12 |
| 3 | Kinurenic acid | 190.0 (+) | 172.0 | 12 |
|  |  |  | 162.0 | 16 |
|  |  |  | 144.0 | 20 |
| 4 | Tryptamine | 161.0 (+) | 144.0 | 12 |
| 5 | Tyramine | 138.0 (+) | 121.1 | 20 |
|  |  |  | 103.0 | 20 |
|  |  |  | 93.1 | 20 |
|  |  |  | 91.0 | 20 |
|  |  |  | 77.0 | 20 |

### Statistical analysis

The experiment was conducted in five biological replicates. All measurements were performed in three replications. To avoid false-positive results indicating the presence of histamine in samples, we employed the methodology that involved analyzing a natural sample, analytical control, and analyzing a sample with a standard control added to it. Mixture of solvents and flushes to rinse Eppendorf tubes and filters was employed as negative control.

The Shapiro-Wilk test revealed that all data sets were normally distributed. The following parametric statistical tests were subsequently applied: prior to conducting a one-way analysis of variance, the uniformity of variance was tested using the Fligner test. (ANOVA using general linear model).

## Results

### Antibacterial and antioxidant activities of V.dubyana extracts

The extracts of laboratory-cultivated *V. dubyana* exhibited weak antimicrobial activity against only two out of eighteen strains of microorganisms involved in the experiment. The moss extracts showed the greatest activity against *St. carnosus* and kanamycin-resistant *E. coli* TolC. It was found that both the methanol extract of *V. dubyana* and the concentrate of the moss environment medium exhibited weak antimicrobial activity against the growth of Gram-positive *St. carnosus*. For the methanol extract of *V. dubyana*, the zone of inhibition of *St. carnosus* was 8 mm. For concentrated water from the aquarium where *V. dubyana* was grown, the zone of inhibition of *St. carnosus* was 6 mm. Also, only methanolic extracts inhibited the growth of kanamycin-resistant *E. coli* TolC, showing the 8 mm inhibition zone. Besides, no fungicidal activity against the tested organisms was observed.

Evaluation of the antioxidant activity revealed moderate qualitative reaction for the methanolic extract of *V. dubyana*, and no qualitative reaction for the moss environment medium.

### Identification of biogenic amines

The UHPLC-MS-MRM analysis detected molecules of tyramine (Fig.1), histamine (Fig.2), tryptamine (Fig.3), kynurenine (Fig.4), kynurenic acid (Fig.5), and in the cell biomass of the moss *V. dubyana*. The analysis of the moss environment medium in aquaria revealed the presence of histamine and the absence of other tested biogenic amines. The quantitative analysis revealed similar concentrations (2±0,3 mg/kg) of histamine both for cell biomass of moss *V. dubyana*, and for the moss environment medium.

**Figure 1.**
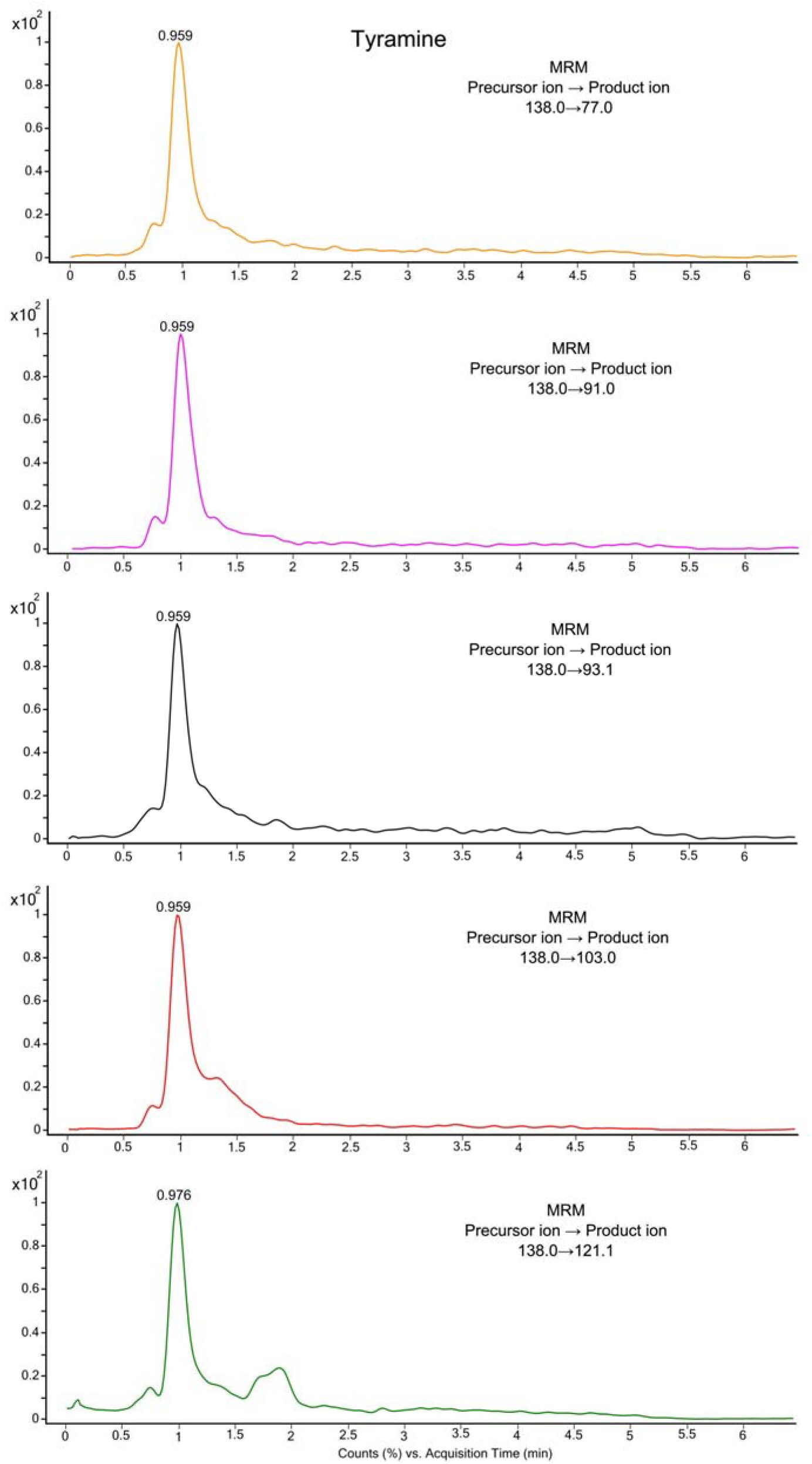
Chromatograms of MRM transitions of tyramine obtained from *V. dubyana* methanol extract

**Figure 2.**
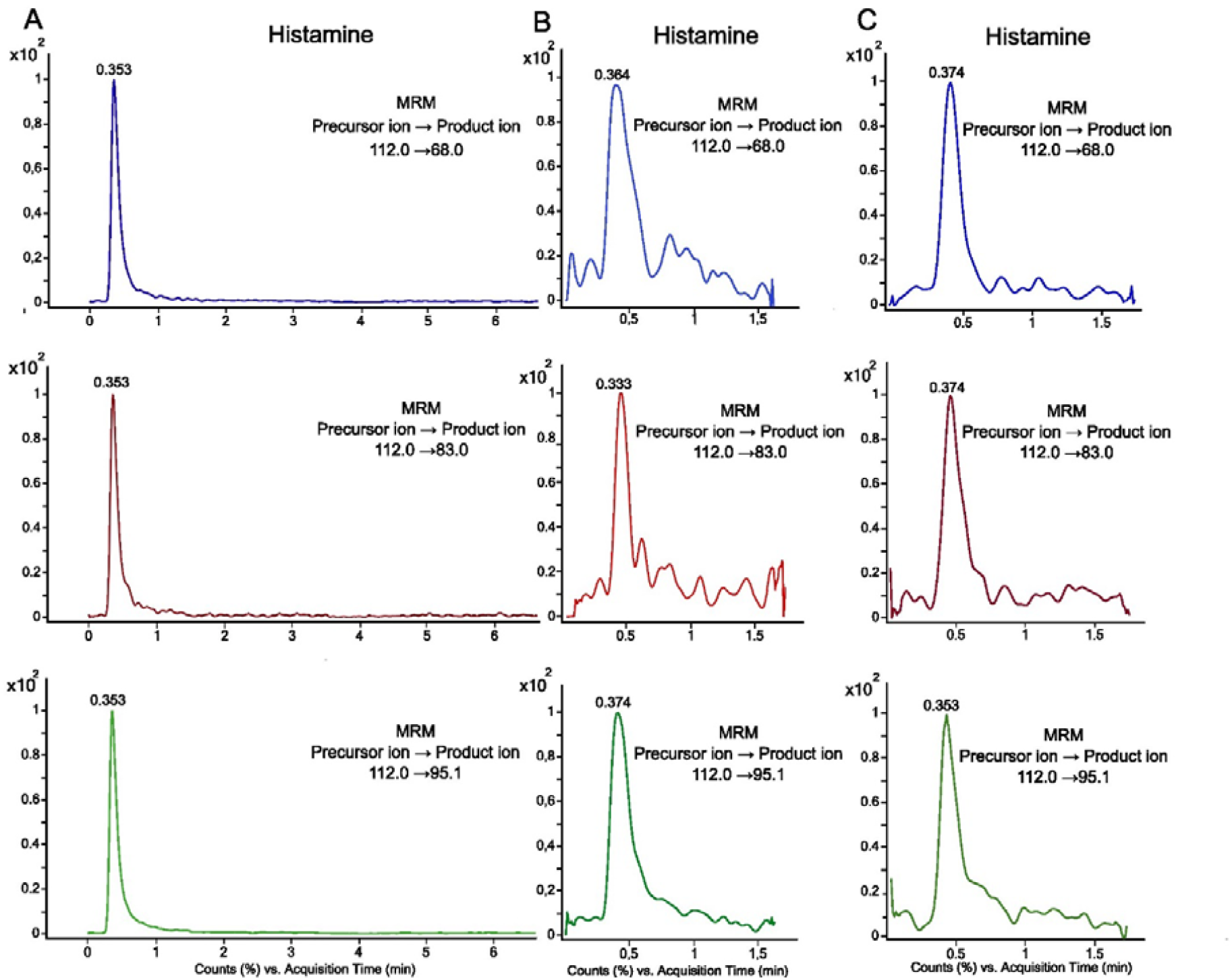
Chromatograms of MRM transitions of histamine: A — chromatogram of histamine dihydrochloride analytical standard (2 mg/L); B — chromatogram of histamine obtained from *V. dubyana* methanol extract; C — chromatogram of histamine obtained from moss environment medium in which *V. dubyana* was grown

**Figure 3.**
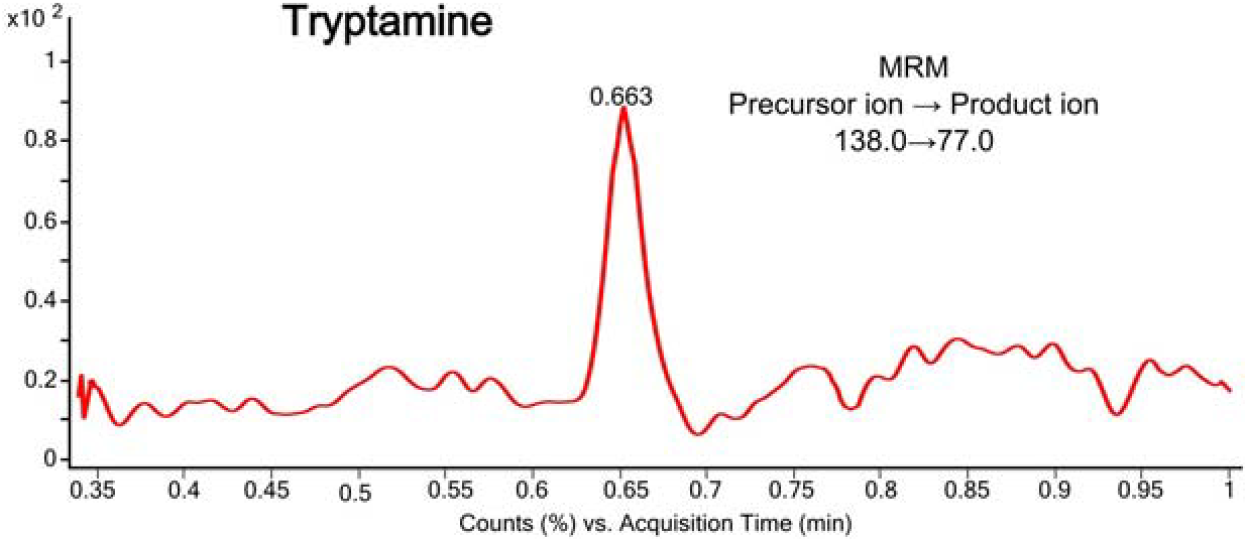
Chromatograms of MRM transitions of tryptamine obtained from *V. dubyana* methanol extract

**Figure 4.**
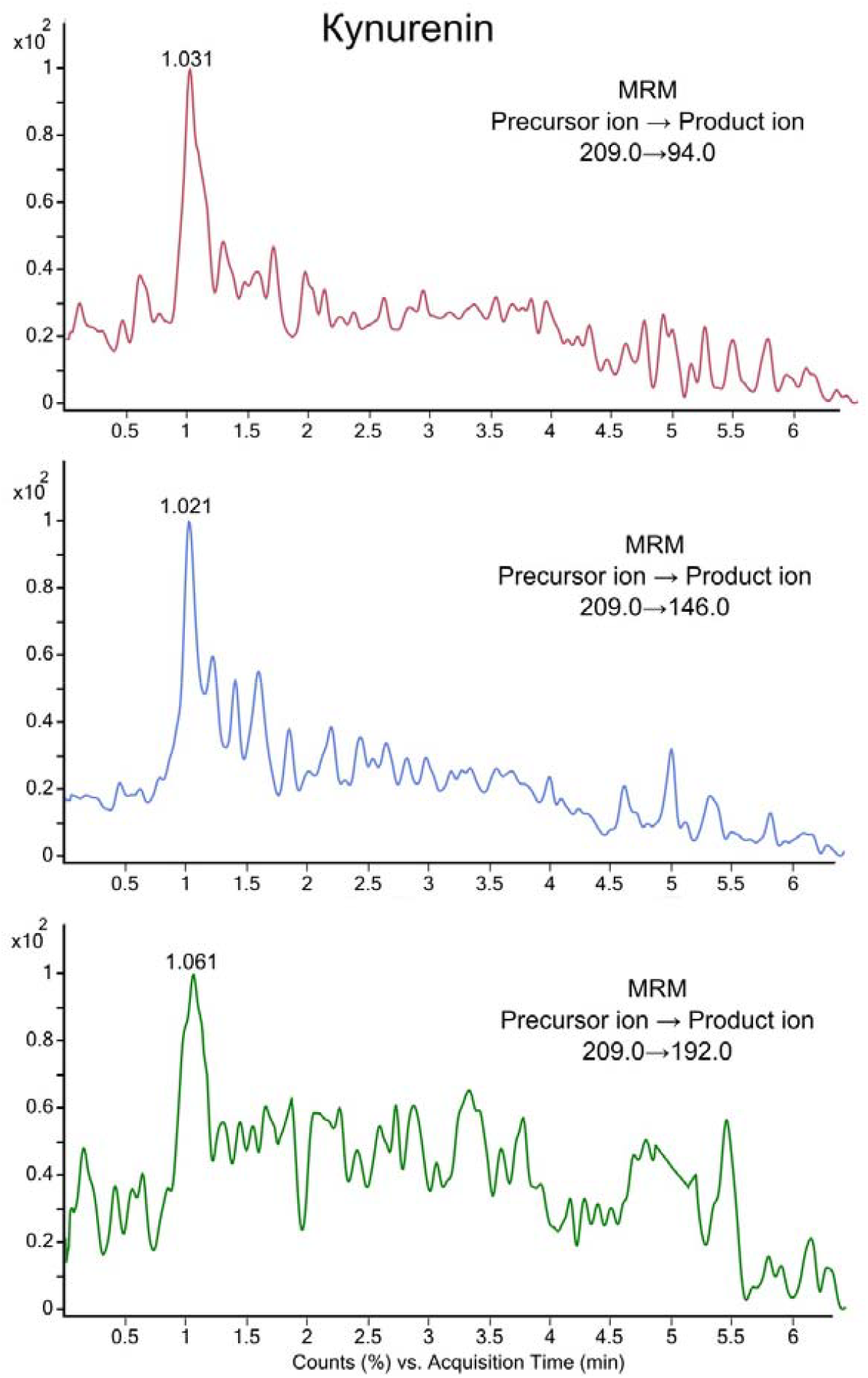
Chromatograms of MRM transitions of kynurenine obtained from *V. dubyana* methanol extract

**Figure 5.**
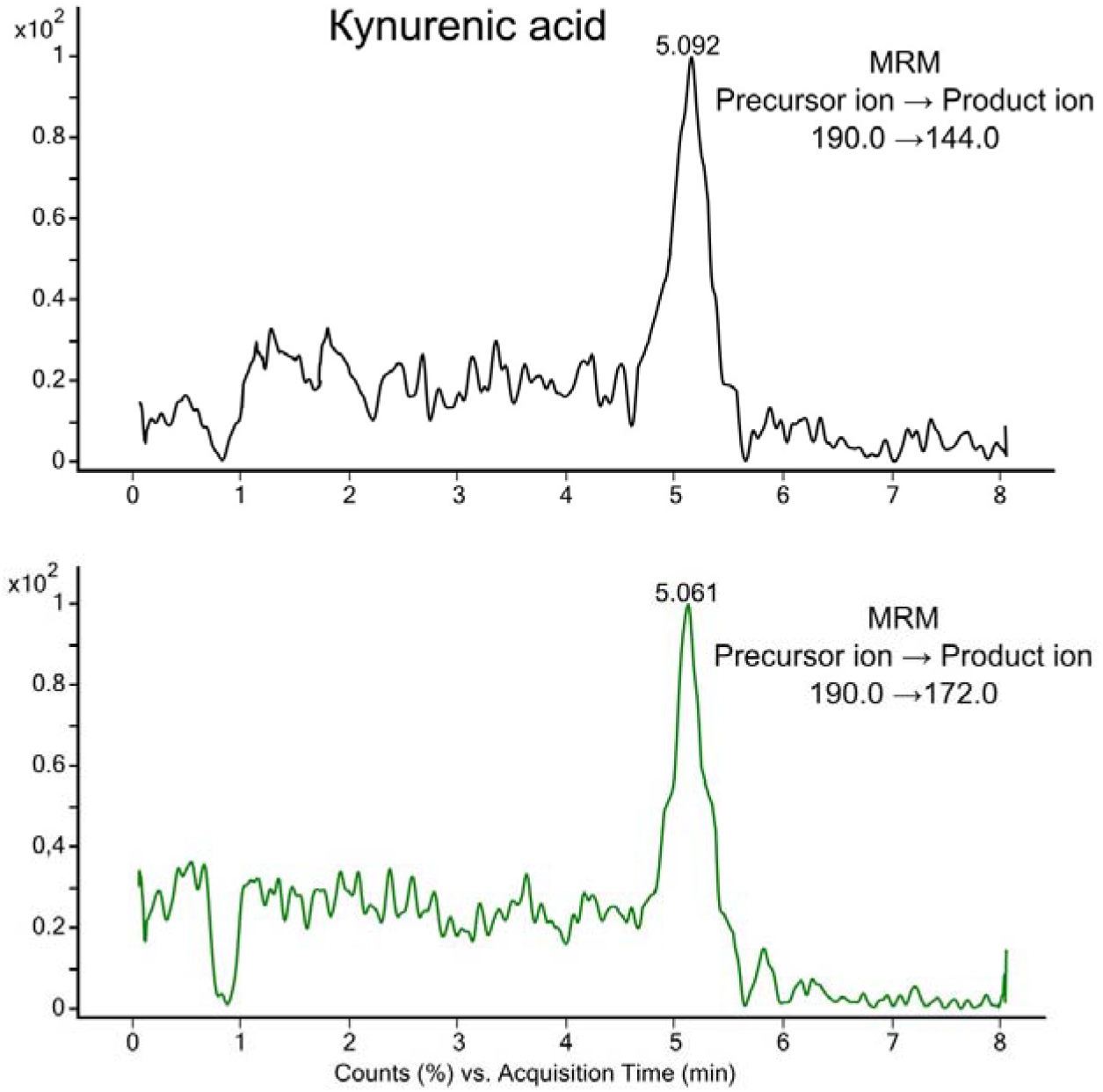
Chromatograms of MRM transitions of kynurenic acid obtained from *V. dubyana* methanol extract

## Discussion

Mankind has always used natural products from plants for a variety of purposes. For example, as food additives, medicines, and perfumes. Because plants produce a wide range of active metabolites, they have always been of great interest to pharmaceutical and chemical industries. For this reason, researchers from different countries focus on synthesis, properties, and evolution of the metabolic diversity of compounds synthesized by plants (Horn et al. 2021; David et al. 2015, Owen et al. 2017).

To date, hundreds of natural compounds of different classes have been isolated from mosses: flavonoids, terpenoids, benzoids, bibenzyls, bis (bibenzyls), phenylpropanoids, and fatty acid derivatives. Extensive studies have been conducted in order to test the properties of extracts and isolated compounds. In addition, the antimicrobial, antiviral, fungicidal, cytotoxic, nematocidal, insecticidal, antioxidant, anti-inflammatory, antifungal, phytotoxic, and other activities of these compounds have been demonstrated (Peters et al. 2019; Dziwak et al. 2022; Horn et al. 2021; Asakawa et al. 2013, 2018, 2022 Sabovljević et al. 2016).

This study is the first to describe the ability of aquatic mosses to produce biogenic amines and natural products that possess antimicrobial and antioxidant properties. The study materials revealed weak antimicrobial and antioxidant activity, which can be explained by the presence of the aforementioned classes of natural compounds that are often responsible for the observed effects.

At the same time, we identify the presence of biogenic amines in the aquatic mosses as the primary result of our study. Biogenic amines serve important functions as hormones, neuromodulators, or neurotransmitters. The term “biogenic amines” was introduced to refer to a group of amines that play a key role in biological processes as chemical intermediaries. They are present in all living organisms including animals, plants, and microorganisms as ancient components (Roshchina 2016). In contrast to animals, cell-to-cell signaling involving biomediators in plants does not occur through gap junctions, but rather through a continuous process. However, properties of these amines have been much less studied in plants than in animals. Monoamine neurotransmitters have been shown to play pivotal roles in plant life, including morphogenesis, flowering, ion permeability, and other processes (Akula and Mukherjee 2020; Roshchina 2016, 2022). It is known that the content of biogenic amines in tissues of higher plants and algae varies from 30 to 700 nMol/g of crude mass under normal conditions, and increases up to 3000 nMol/g of crude mass under stress conditions (Roshchina 2016).

It is worth noting that, until now, studies on aquatic moss species have been relatively limited and mainly referred to their study as effective biomonitoring tools (Carrieri et al. 2022; Real et al. 2021; Świsłowski et al. 2023; Zotina et al. 2024). This study was the first to demonstrate the ability of the aquatic moss *V. dubyana* to produce biogenic amines: histamine, tryptamine, tyramine, kynurenine, and kynurenic acid. To date, we are not ready to explain the role of biogenic amines in aquatic mosses, but we obtained the first data describing the presence of these biochemical mechanisms, which are likely involved in stress adaptation processes in these organisms.

One of interesting and promising findings of our study is the capacity of *V. dubyana* to produce histamine extracellularly. It is known that histamine levels in a plant can vary depending on the state of its development. Some authors have reported increased histamine levels under stress conditions including soil salinity, drought, and parasite infestation (similar response to any stress is also characteristic of animals) (Akula and Mukherjee 2020). The highest concentrations of histamine were identified in the stinging hairs of Urticaceae species (Urtica dioica). The exact location of histamine within the plant cell is not yet known. It is thought that, similar to animal cells, histamine may be concentrated in secretory cells, nuclei, mitochondria, and microsomes. (Roshchina 2016; Akula and Mukherjee 2020). To date, the role of biogenic amines in mosses and, in particular, their cellular locations and metabolic transport systems remains a large and understudied area for future research. Biotechnological use of mosses has gained momentum in recent decades (Reski et al. 2015; Decker and Reski 2020). Despite the relatively extensive interest to biogenic amines in experimental and clinical medicine, actual production of these substances worldwide is relatively limited and basically within the microbiological field. There are several current and historical methods of biogenic amine production. For example, there is a known biotechnological method for producing histamine by bacterial decarboxylation of histidine (Patent SU 166355 A1, “Method of Histamine Production”). Another biotechnological method of histamine production is described in patent WO2013011137A1 “Production and use of bacterial histamine”. This patent describes the method of local histamine biosynthesis by lactic acid bacteria in the gastrointestinal tract of mammals. A chemical process for obtaining tyramine was also described in patent RU 2218326 C2, “Method for obtaining 4-(2-aminoethyl) phenol.” The method allows one to obtain tyramine from available starting compound in two chemical steps with a high yield. To date, no chemical or biotechnological methods for the industrial production of kynurenine and kynurenic acid have been reported. Thus, the studied aquatic moss is a source for simultaneous production of several biogenic amines important for both plants and animals.

It has been demonstrated that the synthesis, storage, degradation and reuptake of biogenic amines are tightly controlled in mammals and play a crucial role in brain function. An imbalance in the levels of these amines can trigger neurodegenerative processes in humans, such as Alzheimer’s disease, Parkinson’s disease, and others (Lindemann and Hoener 2005). Endogenous amine compounds, also known as trace amines, are produced by organisms (plants, bacteria, insects, and mammals including humans) and are expressed in very low concentrations. They include tryptamine, tyramine, and octopamine. It has been reported that the altered levels of trace amines in brain lead to neuropsychiatric disorders such as schizophrenia, attention deficit disorder, depression, and hyperactivity, suggesting that these amines are involved in the pathophysiology of monoaminergic systems (Branchek and Blackburn 2003, Pei et al. 2016).

The use of mosses in biotechnology has several aspects: extracts of mosses or whole plants can be used for various industrial purposes and as a platform for production of valuable metabolites and pharmaceutical proteins. Information on medicinal plants containing neurotransmitters with high biological activity shows the promise of using plant neurotransmitters in pharmaceuticals and biotechnology (Lattarulo 2020; Decker and Reski 2020; Fukuyama 2022).

## Conflict of Interest

The authors declare no conflict of interests.

## Acknowledgements

We thank the Direction of the Biological Faculty and Botanical Garden of Irkutsk State University, and the administration of ISU for their support.

## Funding

The study was carried out with the financial support of the Ministry of Higher Education and Science of the Russian Federation (projects: FZZE 2021-0013, 2024-0003, 2024-0011).

## Notes

### Competing Interest Statement

The authors have declared no competing interest.

